# Assessing the utility of CASP14 models for molecular replacement

**DOI:** 10.1101/2021.06.21.449228

**Authors:** Claudia Millán, Ronan M. Keegan, Joana Pereira, Massimo D. Sammito, Adam J. Simpkin, Airlie J. McCoy, Andrei N. Lupas, Marcus D. Hartmann, Daniel J. Rigden, Randy J. Read

**Affiliations:** Department of Haematology, University of Cambridge, Cambridge Institute for Medical Research, Cambridge CB2 0XY, United Kingdom; Scientific Computing Dept., Science and Technologies Facilities Council, UK Research and Innovation, Didcot, Oxfordshire, United Kingdom; Max Planck Institute for Developmental Biology, Max-Planck-Ring 5, Tübingen, Germany; Institute of Systems, Molecular and Integrative Biology, Biosciences Building, Crown Street, Liverpool L69 7BE, United Kingdom

**Author notes:** These authors contributed equally to the work (Claudia Millán, Ronan M. Keegan). Biozentrum, University of Basel, 4056 Basel, Switzerland (Joana Pereira). <u>Correspondence to</u>: Randy J. Read, Department of Haematology, University of Cambridge, Cambridge Institute for Medical Research, The Keith Peters Building, Hills Road, Cambridge CB2 0XY, U.K.

**Keywords:** structure prediction, model refinement, CASP, molecular replacement, likelihood

## Abstract

The assessment of CASP models for utility in molecular replacement is a measure of their use in a valuable real-world application. In CASP7, the metric for molecular replacement assessment involved full likelihood-based molecular replacement searches; however, this restricted the assessable targets to crystal structures with only one copy of the target in the asymmetric unit, and to those where the search found the correct pose. In CASP10, full molecular replacement searches were replaced by likelihood-based rigid-body refinement of models superimposed on the target using the LGA algorithm, with the metric being the refined likelihood (LLG) score. This enabled multi-copy targets and very poor models to be evaluated, but a significant further issue remained: the requirement of diffraction data for assessment. We introduce here the relative-expected-LLG (reLLG), which is independent of diffraction data. This reLLG is also independent of any crystal form, and can be calculated regardless of the source of the target, be it X-ray, NMR or cryo-EM. We calibrate the reLLG against the LLG for targets in CASP14, showing that it is a robust measure of both model and group ranking. Like the LLG, the reLLG shows that accurate coordinate error estimates add substantial value to predicted models. We find that refinement by CASP groups can often convert an inadequate initial model into a successful MR search model. Consistent with findings from others, we show that the AlphaFold2 models are sufficiently good, and reliably so, to surpass other current model generation strategies for attempting molecular replacement phasing.

## Introduction

As protein structure prediction becomes more accurate and reliable, it is becoming an increasingly useful tool in a variety of scenarios, such as prediction of the structural context of mutations either associated with disease or with escape from an immune response. It is also clear that protein structure prediction will accelerate the experimental determination of 3D structures, by improving the models for molecular replacement (MR).

MR is the most commonly used method to determine the unmeasured phases needed to compute an electron density map from a diffraction pattern. This is carried out, typically, by determining the orientation angles and translation vector (together referred to as the ‘pose’) required to superimpose the model generated by prediction with the coordinates of the atoms in the crystal. Models generated by structure prediction supplement the models that can be derived from previously-determined structures of homologues in the worldwide Protein Data Bank (wwPDB)^1^, often involving extensive editing.

As recently as 20 years ago, it would have been fair to say that even template-based protein models were rarely more useful for MR than the templates on which they were based, because it was too difficult to distinguish the few ways in which they could be improved from the vast number of ways in which they could be degraded. Since then, modeling methods have turned a corner and are becoming progressively more useful. A test for utility in MR was introduced for CASP7^2^, showing that about half of the best available templates in the high accuracy category could be improved by at least one predictor group, although only 33 of 1588 models evaluated were better than the best template. It should be acknowledged here that there is less room for improvement in the high accuracy category than in cases where no closely related template is available. Indeed, in a striking case from CASP7, an *ab initio* model of a small globular protein was predicted to sufficient accuracy that it could have been used to solve that structure by MR^3^. Other work resulted in the program AMPLE which seeks to isolate sufficiently accurate substructures from sets of *ab initio* models by clustering and truncation^4^.

When model accuracy was low, a useful score could only be generated if the model was sufficiently good to identify the correct pose in the full search. This problem was circumvented later by the use of rigid-body refinement starting from a structural superposition instead of the full MR search, which also had the benefit of dramatically reducing the CPU time required to explore many incorrect solutions with poor models that lack useful signal, and ensuring that the LLG scores corresponded with models in the correct pose. Although the success-or-failure aspect of the MR searches was lost, the LLG scores could still be interpreted in the knowledge that MR searches yielding LLG values above 60 are usually correct^5^.

A second problem arose in MR scoring when there are multiple copies in the asymmetric unit, or more than one type of component. With the full MR approach, the MR scoring was restricted to those cases for which there was a single copy of a single protein component in the asymmetric unit of the crystal. However, the rigid body refinement approach allowed these more complicated targets to be scored by placing all copies of the tested model within a background that includes the deposited structure for all other components of the crystal; the increase in the LLG obtained when adding the tested model to the background structure alone was the measure of model quality.

A *Phaser* script to carry out rigid-body refinement approach calculations, written by Gábor Bunkóczi, was used by other assessors in the refinement category of CASP10^6^, as well as by us for both the refinement^7^ and template-based modeling^8^ categories of CASP13. This script was again used here for assessment in CASP14.

Problems remain with the rigid-body refinement approach, not least the fact that it requires diffraction data to be made available to assessors; not all crystallographers contributing targets are able to share these data in advance of publication. A substantial number of targets and domain evaluation units (EUs) derived from them now arise from cryo-EM structure determinations (21 EUs from 7 structures in CASP13^9^, and 22 EUs from 7 structures in CASP14) and hence have no diffraction data. In addition, the LLG scores vary in a crystal-form-dependent fashion, depending on the resolution and quality of the data, the number of copies of the protein in the asymmetric unit of the crystal, and the fraction of the asymmetric unit accounted for by the modeled component. Comparisons among targets require some normalization, generally through the calculation of Z-scores.

In this work, a novel likelihood score is introduced, the ‘relative expected LLG’ (reLLG) that requires only the coordinates of the target to rank the suitability of a model for MR. Most significantly, it is a crystal-form independent measure. We test the reLLG against the LLG score as a ranking measure and demonstrate its utility as a more convenient and robust measure, which should supersede the use of the LLG for this purpose. We find that the ability of refinement groups to improve reLLG values correlates well with their ability to improve the performance of refinement targets in actual MR experiments. Finally, our results provide another metric by which the superiority of the AlphaFold2 (Jumper *et al.,* this volume) models over the others in the assessment can be seen.

## Materials and Methods

### Target selection for log-likelihood-gain scoring

In CASP, structures contributed for the prediction season are examined and divided into smaller pieces (often individual domains) that usually have a relatively compact structure. These are referred to as “evaluation units” or EUs. For CASP14, a total of 96 EUs were selected for evaluation of structure prediction. Prior to the CASP14 meeting, diffraction data were made available by the experimentalists who contributed 32 crystal structures, from which 54 EUs were drawn. These EUs were therefore able to be included in the MR assessment, which used the previously described diffraction data-dependent LLG score. Diffraction data were not available at the time of assessment for the remaining 17 EUs drawn from other crystal structures, nor of course for the EUs drawn from cryo-EM or NMR structures.

In the refinement round, a total of 49 prediction targets were selected. These included 7 “extended” targets and 7 “double-barrelled” targets used to conduct additional experiments in CASP14. For the extended targets, refined models were collected after the initial 3-week period and again after an additional 6 weeks, during which more extensive computations may have been performed (denoted with an “x” in the target name). For the double-barrelled targets, two starting models were chosen for refinement, one typically chosen from the server models and the other from models submitted by the AlphaFold2 group (denoted with “v1” or “v2” in the target name, with the “1” or “2” chosen randomly). Thirty-four of the 49 total targets were derived from structures determined by X-ray crystallography, of which 20 had diffraction data available at the time of assessment and could therefore be used for LLG calculations.

### Model selection

For the double-barrelled refinement targets, one group recognized correctly that one of the two starting models (the AlphaFold2 model, though it was not identified as such) was superior to the other, and they submitted the better model as a refinement model for the poorer one. While the ability to recognize good models is laudable, it does not reveal anything about the ability of the group to carry out refinement, so the Alphafold2 models provided by this group were excluded from consideration. All other models for both structure prediction and refinement were evaluated.

### Evaluation measures

#### Log-likelihood-gain

As in the case of CASP13, the LLG for each model of each EU was computed by rigid-body refinement in *Phaser,* using the rest of the final crystal structure as a fixed background for the calculation. The initial superposition of the evaluation unit on the target was carried out using the sequence-independent structure alignment program TM-align^10^. To allow for an assessment of the impact of the predicted error estimates, the LLG calculations were performed in two different modes for each prediction: once with the B-factor field interpreted as error estimates (used to weight the MR calculations as discussed below) and once with all B-factors set to a constant value. From each of these scores, we subtracted the EU-specific null-model LLG (the LLG value of the models with the lowest GDT_HA, corresponding to the noise), thus calculating the equivalent to the CASP13 increase in LLG from the background. The definition of the EU-specific null-model-LLG stems from the observation that at low GDT_HA, LLG values in GDT_HA vs LLG plots can be approximated by linear regression for a given EU.

To calculate the EU-specific null-model-LLG, for each EU, the models were binned into 100 equally spaced GDT_HA bins and the average LLG value for each bin taken. This average was computed iteratively, removing at each iteration those data points with an LLG 1σ below the average until no more data points were excluded. Out of these bins, the first 35% (bottom 35% GDT_HA) were considered further, and the average of their average LLG taken. Those bins with an average LLG within 3σ the average over all bottom 35% were sorted by their average LLG and the middle 80% taken. A linear model was fitted to the averages of these bins and the intersection in the *y* axis taken as the null-model-LLG. All models with an LLG below the corresponding null-model-LLG were assigned a score of zero. This can happen if the entries in the B-factor field for a model are all too large to correspond to sensible root-mean-square displacement (RMSD) estimates and effectively downweight the contributions of the atoms to zero. The likelihood calculations can also fail for computational reasons, such as if the model represents an unfolded protein and extends over such a large volume that memory limits are exceeded in the FFT calculations of structure factors. Models leading to such failures are also assigned a score of zero.

We refer to the difference LLG score as the dLLG for short.

#### Relative expected log-likelihood-gain

As discussed above, there are substantial advantages to a likelihood score that measures suitability for MR independent of crystal form or structure determination method.

By the correlation theorem of Fourier transforms, the correlation between electron densities is proportional to the complex correlation between structure factors calculated from those electron densities. In turn, the complex correlation in a resolution shell is equivalent to the resolution-dependent *σ_A_* value used in crystallographic likelihood targets, such as the log-likelihood gain on intensities (LLGI) used for MR^11^. (Note that the complex correlation in a resolution shell is also equivalent to the Fourier shell correlation, or FSC, commonly used to assess cryo-EM reconstructions^12^.) We have shown that there is a close relationship between *σ_A_* and the score expected to be obtained in likelihood-based MR. The expected log-likelihood-gain (eLLG) can be approximated^5^ as the sum, over all Fourier terms, of 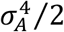, allowing valuable optimizations of the MR strategy depending on the qualities of the model and the data^13^. This relationship between electron density overlap and LLG is the basis of the reLLG score discussed below.

Superposition of model and target with an algorithm such as that in the LGA program^14^ will not generally optimize the electron density overlap. Therefore, to enable the calculation of the reLLG score, a new phased rigid-body refinement mode was implemented in *phasertng*, which is under development to replace and enhance the functionality of *Phaser*^15^. The rigid-body refinement starts from a sequence-independent superposition using LGA^14^. Instead of optimizing the LLGI score, which lacks phase information, it uses a phased likelihood target. This target starts from the assumption, based on the Central Limit Theorem, that structure factors computed from two superimposed models are related by a bivariate complex normal distribution; the assumption of multivariate complex normal distributions also underlies many likelihood-based crystallographic algorithms, including MR, refinement and experimental phasing. The probability distribution relating two sets of structure factors is characterized by a Hermitian covariance matrix. This takes a particularly simple form if the structure factors are first normalized, giving E-values for which the mean-squared value is one. In this case, the off-diagonal complex covariance term of the covariance matrix becomes the complex correlation, *σ_A_*:

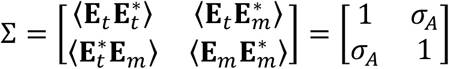

Note that a complex covariance will in general be a complex number, but *σ_A_* is a real number because, if a systematic phase shift were known between the two structures, that would imply the existence of a known relative translation vector, which could be corrected instead.

The likelihood target is the conditional probability of the target structure factors given the known model structure factors. This is derived from the joint distribution by standard manipulations to obtain the conditional variance of the target E-value given the model and its expected value:

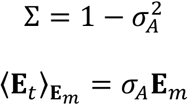

These parameters are used to express the conditional probability as a complex normal distribution:

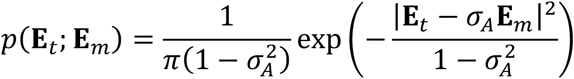

The target that is optimized is a log-likelihood-gain, obtained by taking the logarithm of the conditional probability and subtracting the logarithm of the probability of the null hypothesis, which is the Wilson distribution of structure factors^16^ and is equivalent to the conditional probability when *σ_A_* is zero, *i.e.* when the model is uncorrelated to the target and is thus uninformative. The contribution of a single Fourier term to the total LLG is given in the following:

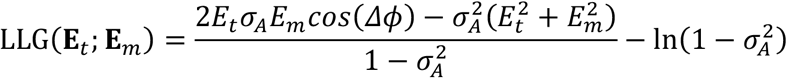

The phased log-likelihood-gain is a function of the orientation and position of the model relative to the target, and of the current value for *σ_A_* for each structure factor pair. The orientation is defined in terms of three rotation angles specifying rotations of the preoriented model around axes parallel to *x, y* and *z* running through the center of the model. Because the perturbations of the initial orientation will be small, these rotations will be nearly orthogonal and will therefore behave well in the optimization. The position is defined in terms of translations along the *x*, *y* and *z* axes, which are orthogonal and are essentially independent of the rotations applied around the center of the model. The *σ_A_* values are a function of the resolution of the relevant structure factors and are defined in terms of the radial RMSD for coordinate errors drawn from a single 3D Gaussian. The value of *σ_A_* is given, as a function of resolution, by the Fourier transform of that Gaussian:

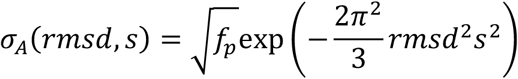

where *f_p_* is the fraction of the target explained by the model, assumed to be one for the calculations reported here, *rmsd* is the refined parameter and *s* is the magnitude of the diffraction vector (the inverse of the resolution).

The refinement against the phased log-likelihood-gain can be seen to optimize the electron density overlap: *E_t_E_m_cos*(*Δϕ*) is equivalent, by the correlation theorem, to the contribution of a Fourier term to the density correlation. The variance term in the phased log-likelihood-gain is controlled by the *rmsd* parameter, which will be optimal when the *σ_A_* values computed as a function of resolution from that *rmsd* match the mean values of *E_t_E_m_cos*(*Δϕ*) in resolution shells.

Once an optimal superposition is obtained, structure factors from the target and the superimposed model are compared in resolution shells to obtain an FSC curve. The eLLG is then calculated by accumulating the sum of *FSC*^4^/2, weighted by the number of Fourier terms in each shell. This eLLG varies with the number of Fourier terms, determined by crystal lattice volume (size of the target), so normalization to a relative eLLG (reLLG) is required to put the scores on a common scale for all target-model pairs. The normalization cannot be carried out simply by comparing the eLLG to what one would expect for a perfect model, because the conditional probability for a perfect model with perfect data is a delta function, which would yield an infinite LLG. We resolve this problem by introducing an “ideally imperfect model”, that is, the best model one expects to get from a high-resolution structure determination, given the limitations of a static model in portraying an ensemble of conformations and the differing influence of crystal packing and other environmental effects. By comparing structures of the same protein in different crystal forms^17^ and by extrapolating the dependence of structural variation with sequence identity to 100% identity^18^ it emerges that the best one might hope for is an effective *rmsd* of about 0.4 Å. The reLLG is therefore computed by dividing the eLLG for the model being tested by the eLLG that would be obtained for *σ_A_* values computed for a complete model with *rmsd* set to 0.4 Å.

The reLLG calculation also requires making a choice for the high resolution limit. A calculation carried out to a higher resolution limit would be more sensitive to model errors, whereas the use of lower resolution would be more forgiving. In principle, one could define scores based on different resolution limits, analogous to the way that the GDT_TS score is more forgiving than the GDT_HA score^14^. We have chosen a resolution limit of 2 Å for calculations here for two reasons. First, the median resolution of crystal structures in the wwPDB^1^ is close to this value: 2.2 Å overall, and 2.1 Å for the year 2020. Second, 2 Å is approximately the resolution at which most structures can be completed starting from even a partial correct MR solution^19^.

We note that it would be possible to compute an eLLG from the *σ_A_* curve defined by the refined *rmsd* parameter, and this could even be done analytically. The advantage of using the actual FSC curve from the comparison of structure factors is that no assumptions are made about the distribution of coordinate errors in the model. The use of a single *rmsd* requires that all the coordinate errors are drawn from the same 3D Gaussian distribution, whereas models have locally varying errors. It is further assumed that the coordinate errors and the atomic scattering factors are uncorrelated, whereas atoms on the surface of a protein both tend to have higher B-factors and are modeled less accurately^20^.

#### Measuring the utility of coordinate error estimates

For a number of years, predictors submitting models for CASP have been asked to provide estimated RMS positional errors in the B-factor field of the PDB files containing the models, on the principle that knowing how confident you should be in a model is as useful, in practice, as the model itself. By CASP13, most predictors in the template-based modeling category included error estimates^8^ but many participants in the refinement category did not^7^. In this round of CASP, we were pleased to see that most predictors and participants in the refinement category do seem to have provided coordinate error estimates within a plausible range.

Such error estimates are extremely valuable for MR models. If the B-factors of the models are increased by an amount that effectively smears each atom’s density over its probability distribution of true positions using the following equation, the electron density overlap, and therefore the LLG score, is optimized.

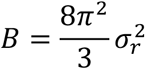

This approach was suggested in the high-accuracy assessment for CASP7^2^ and supported by tests using either the actual or estimated coordinate errors in models^21^. The practical impact was demonstrated further by showing that this treatment significantly improves the utility for MR of models submitted to CASP10^22^, as well as in the evaluation of template-based modeling for CASP13^8^.

To measure the utility of the error predictions numerically, each model was evaluated two times. In the primary calculation, the number in the B-factor field of the model was transformed using the equation above from a coordinate error estimate into a B-factor providing an error weight; in the secondary calculation the B-factor was substituted with a constant value set to 25 Å^2^. (Because the calculation is carried out with normalized structure factors, or E-values, the actual value of the constant B-factor is irrelevant. By extension, the mean value of any B-factor distribution can be altered without affecting the result.) The difference between the two results is a measure of the value added by the error estimates.

### Computing group rankings

For all the evaluation measures, Z-scores were computed using an algorithm that has frequently been applied in other rounds of CASP. The primary ranking was based on model #1 of up to 5 models submitted for each target; this choice implicitly rewards the ability of groups to assess the relative quality of their models. Z-scores were computed in two steps: a set of initial scores was calculated based on the mean and standard deviation (SD) of all models under consideration. All models yielding a Z-score below −2 in the first pass were considered as outliers and the Z-scores recomputed using the mean and SD obtained when the outliers were excluded. At the end, the minimum Z-score was set to −2 to avoid excessively penalizing outliers. For ranking, all Z-scores were summed and a penalty of −2 introduced per target for which a method did not produce a model, effectively treating missing models as outliers.

For rankings based on either the conventional LLG or the new reLLG score, the primary ranking was based on interpreting the B-factor field as an estimate of the RMS error in that atomic position, as requested in the submission instructions provided by the CASP organizers. The difference between this LLG or reLLG for error-weighted models and the value computed setting all B-factors to a constant value was used to measure the value added by the coordinate error estimate.

### Software and data availability

The tables with the reLLG calculations as well as the Jupyter notebooks^23^ used to analyze them can be found in the following repository: https://github.com/clacri/CASP14_MR_evaluation. The Jupyter notebooks have been prepared to be run in the cloud environment of Google Collaboratory^24^, so that the results can be reproduced without having to set up a specific local environment. The analysis relies on the following python scientific libraries: Matplotlib^25^, Pandas^26^, Numpy^27^.

Computation of the reLLG was implemented in *phaser_voyager* (manuscript in preparation), a structural biology computing framework that exploits *phasertng*^15^ in its core. Its focus on modularity and abstraction enables rapid implementation of specific strategies, tracking of pathways, and result analysis. The *phaser_voyager* strategy ‘CASP_rel_eLLG’ will be released as part of version 2 of Phenix^28^, available from the command line as phenix.voyager.CASP_rel_eLLG, requiring the pdb of the target structure and a path to the folder containing approximately oriented models to evaluate.

## Results

### Structure prediction assessment

The statistical analysis and ranking calculations were carried out as described in Materials and Methods. Briefly, the primary ranking was based on the sum of the Z-scores for the #1 predictions when the B-factor field was interpreted as an error estimate, and including the penalty of assigning a Z-score of −2 for missing models.

#### Group rankings by difference log-likelihood-gain (dLLG) scores

Conventional dLLG scores were calculated for 54 evaluation units that correspond to the 32 targets for which the experimental diffraction data were available to us at the time of assessment. We calculated the scores with and without using the error estimates that were intended to be encoded in the B-factor field, thus assessing the impact of the error estimates. The resulting rankings are shown in Fig. 1.

**Figure 1.**
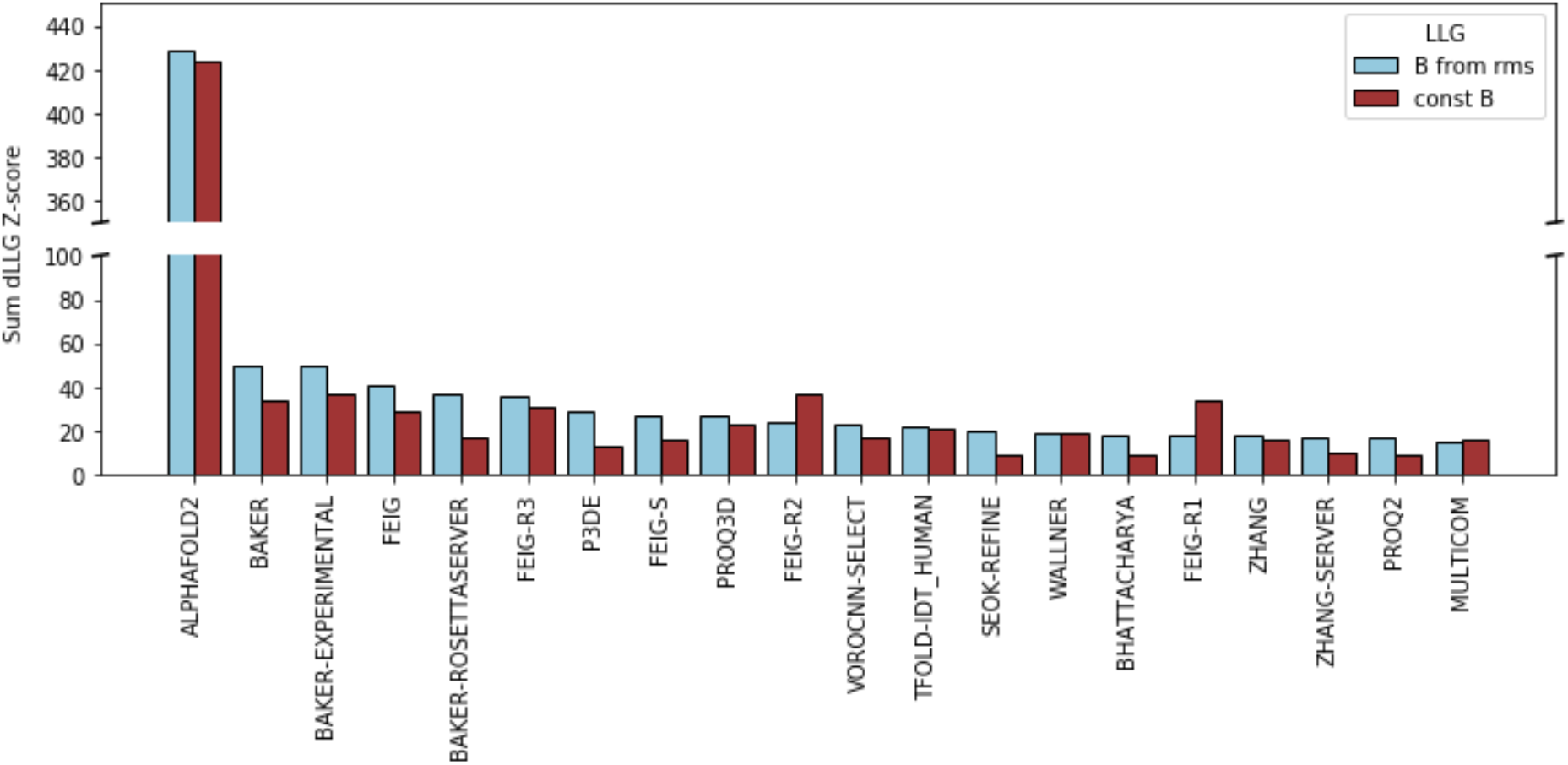
The top 20 groups ranked by the sum of Z-scores of the dLLGs for their #1 predictions. Methods were ranked based on the dLLGs computed when considering the values in the B-factor field as error estimates (predicted RMSD to the target).

#### Group rankings by reLLG scores

146 groups presented models for at least some of the 96 EUs. While calculation of the dLLG score requires diffraction data (limited at the time of assessment to 54 EUs), the calculation of the reLLG does not, and so allows all 96 EUs to be included in the statistics. This includes 71 EUs derived from structures determined by X-ray crystallography, 22 from cryo-EM structures and 3 from NMR structures.

In order to compare and assess the novel reLLG score against the traditional CASP dLLG score, we addressed three questions. First, do the dLLG and reLLG yield similar rankings of models for a specific target? Second, do the dLLG and reLLG yield similar group rankings? Third, do the reLLG calculations obtained from cryo-EM or NMR experiments also yield correlated group rankings?

We compared reLLG scores with dLLG scores for the targets for which diffraction data were available at the time of assessment. We do not expect these measures to be linearly related to each other, because the dLLG score is affected non-linearly by factors such as model quality (which has different effects for different resolution limits) or the fraction of the asymmetric unit of the crystal accounted for by the model. Because the reLLG calculation has been designed to cope better with numerical issues caused by the large estimated RMS errors found in some CASP models, comparisons of the scores obtained interpreting the B-factor field as an estimated error can be complicated by the relative instability of the *Phaser* calculations with some models. To avoid these complications, we chose to compare the reLLG and dLLG values obtained when setting the B-factors constant. Fig. 2 shows scatter plots for four disparate cases, spanning different degrees of modeling difficulty, different fractions of the asymmetric unit accounted for by the model, and different resolution limits. The relationship between the two scores is roughly monotonic, indicating that they will deliver similar ranking order for models.

**Figure 2.**
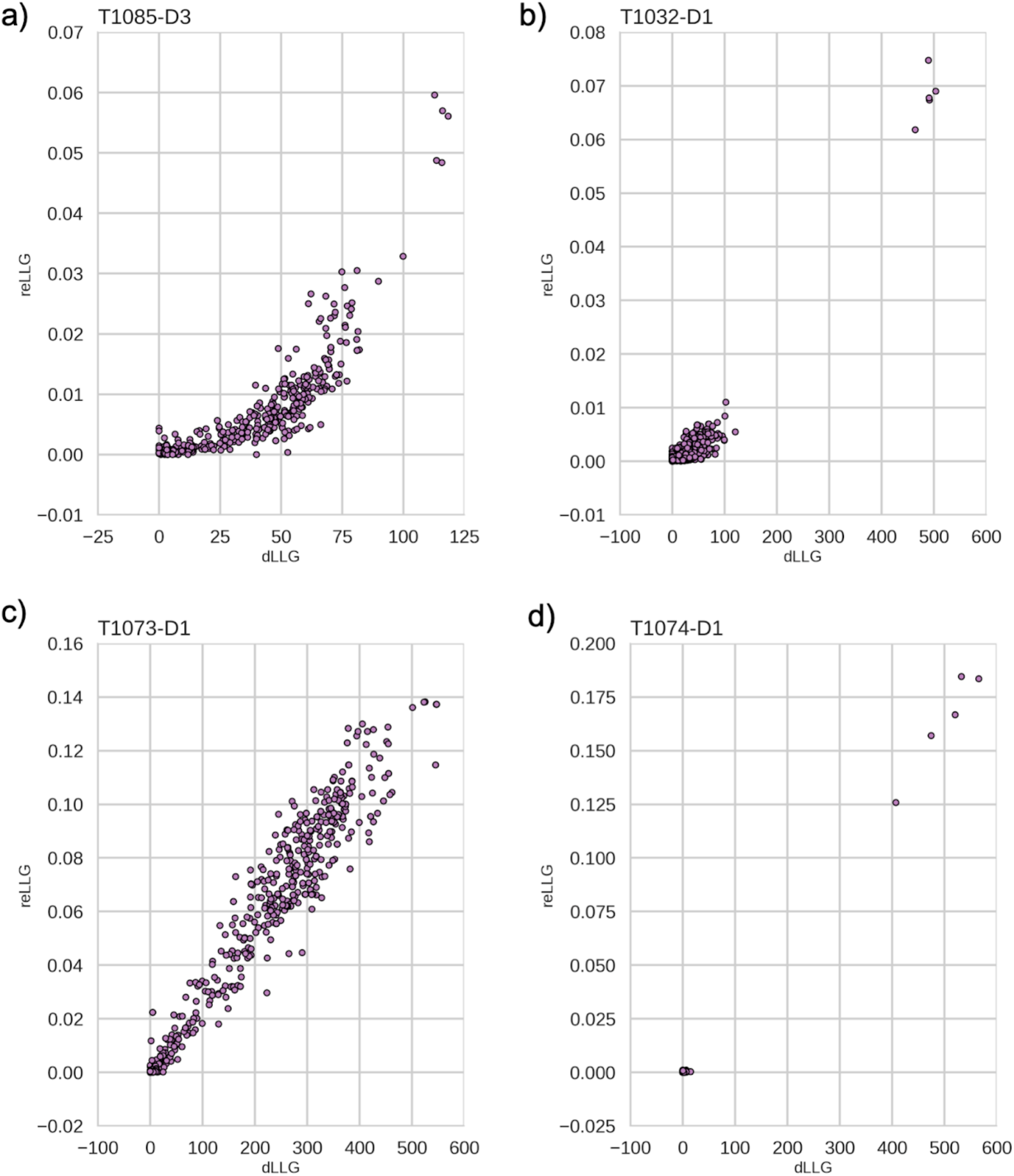
Scatter plots comparing dLLG and reLLG scores for models of 4 EUs illustrating different circumstances. a) T1085-D3: TBM-hard case in which the model comprises 9.8% of the structure, data to 2.26 Å resolution. b) T1032-D1: TBM-hard case, 6 copies of the target in the asymmetric unit, data to 3.3 Å resolution. c) TBM-easy case, 2 copies, data to 1.9 Å resolution. d) FM case, 1 copy, data to 1.5 Å resolution.

Next, we examined whether the group ranking on the subset of targets for which diffraction data were available was similar. Fig. 3 shows a very strong correlation between the ranking orders, with the top 5 groups being identical for the two measures.

**Figure 3.**
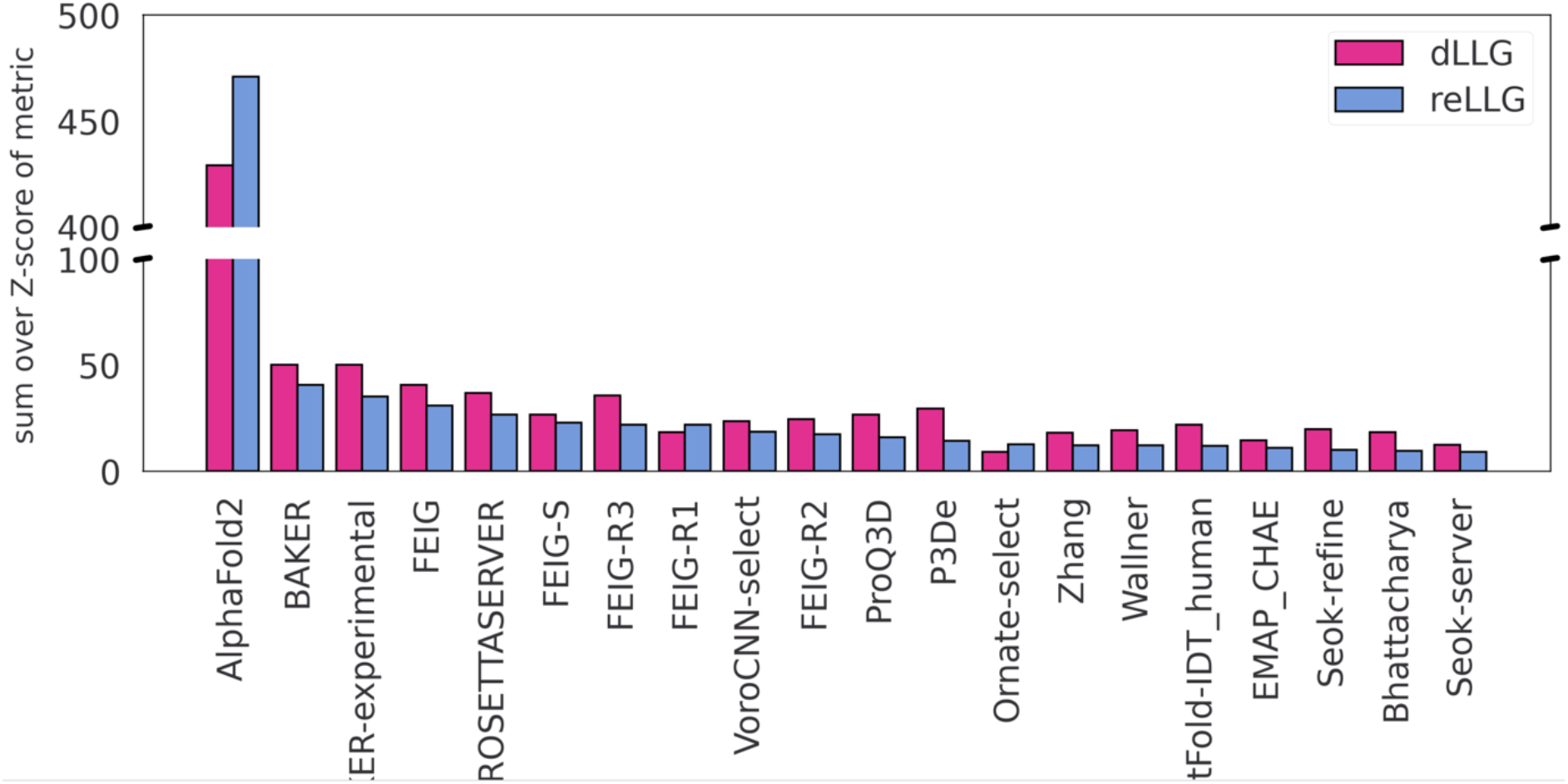
Ranking scores based on dLLG (magenta bars) and reLLG (blue bars) using only targets for which diffraction data were available at the time of assessment. Groups are ordered by their reLLG ranking score.

To verify that there are no systematic differences in how reLLG would score models of structures determined by other methods, we compared the group ranking scores that would have been achieved using only cryo-EM targets or NMR targets with those achieved using X-ray targets. The scatter plots in Fig. 4 demonstrate a strong correlation among the rankings using all three types of target. Note that the NMR scores are based on only 3 EUs.

**Figure 4.**
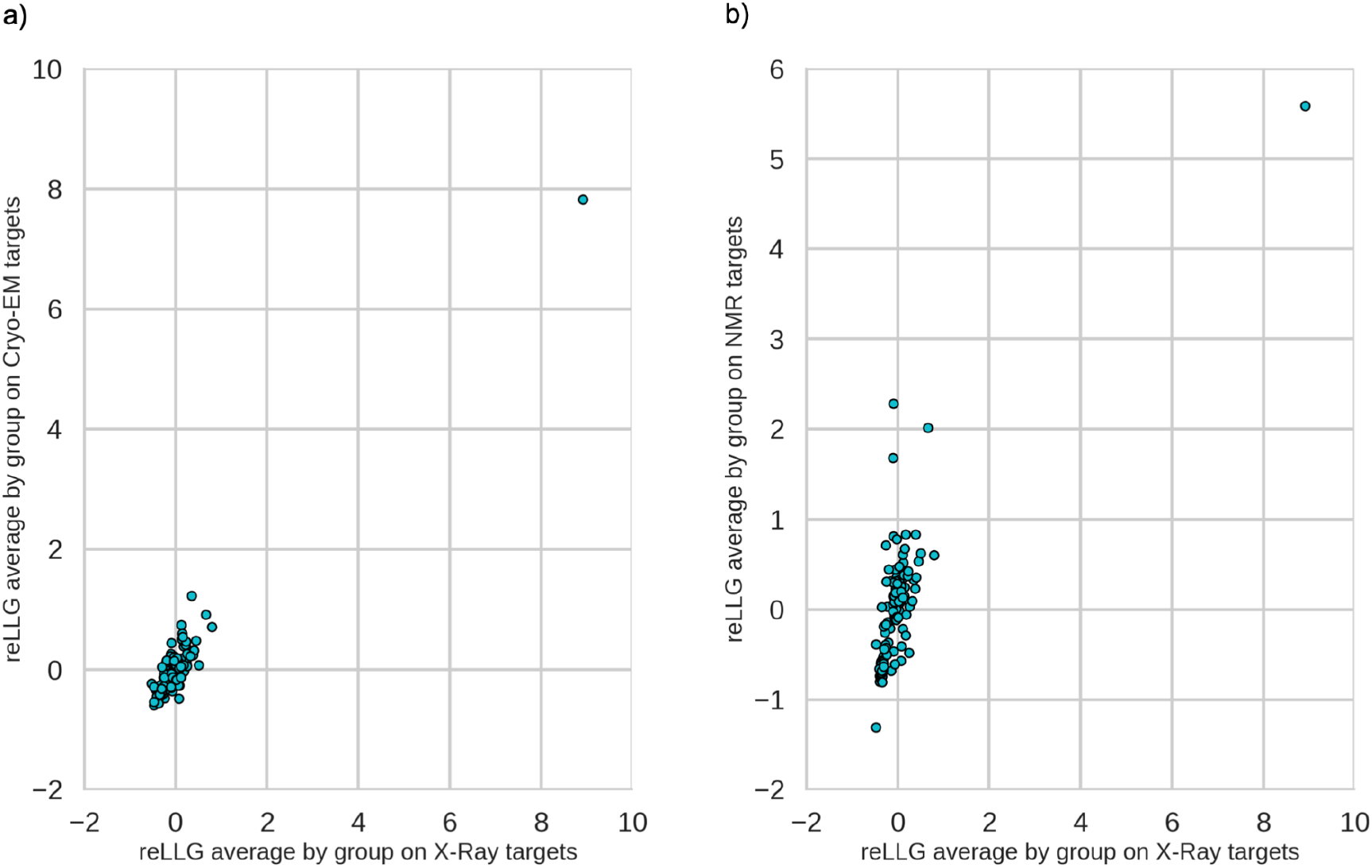
Scatter plots comparing average reLLG scores per group by experimental technique. a) X-ray vs CryoEM. b) X-ray vs NMR.

Given that rankings on common targets are very similar using either dLLG or reLLG, that reLLG rankings on sets of targets derived by different methods (X-ray, cryo-EM, NMR) are similar, and that the use of the reLLG allows the use of a much larger data set (96 EUs rather than 54), we expect the ranking based on reLLG to be closer to what would be achieved for dLLG if diffraction data were available for all 96 EUs than the dLLG ranking based on only 54 EUs. The ranking based on reLLG is more robust, and we take it as the authoritative ranking for this study.

The ranking for all targets by reLLG Z-score (Fig. 5a) is again dominated by AlphaFold2, as also seen with dLLG and more traditional CASP measures. The other top performing groups are BAKER, BAKER-experimental, FEIG2 and BAKER-ROSETTAserver, followed by a few other variants of FEIG group algorithms.

**Figure 5.**
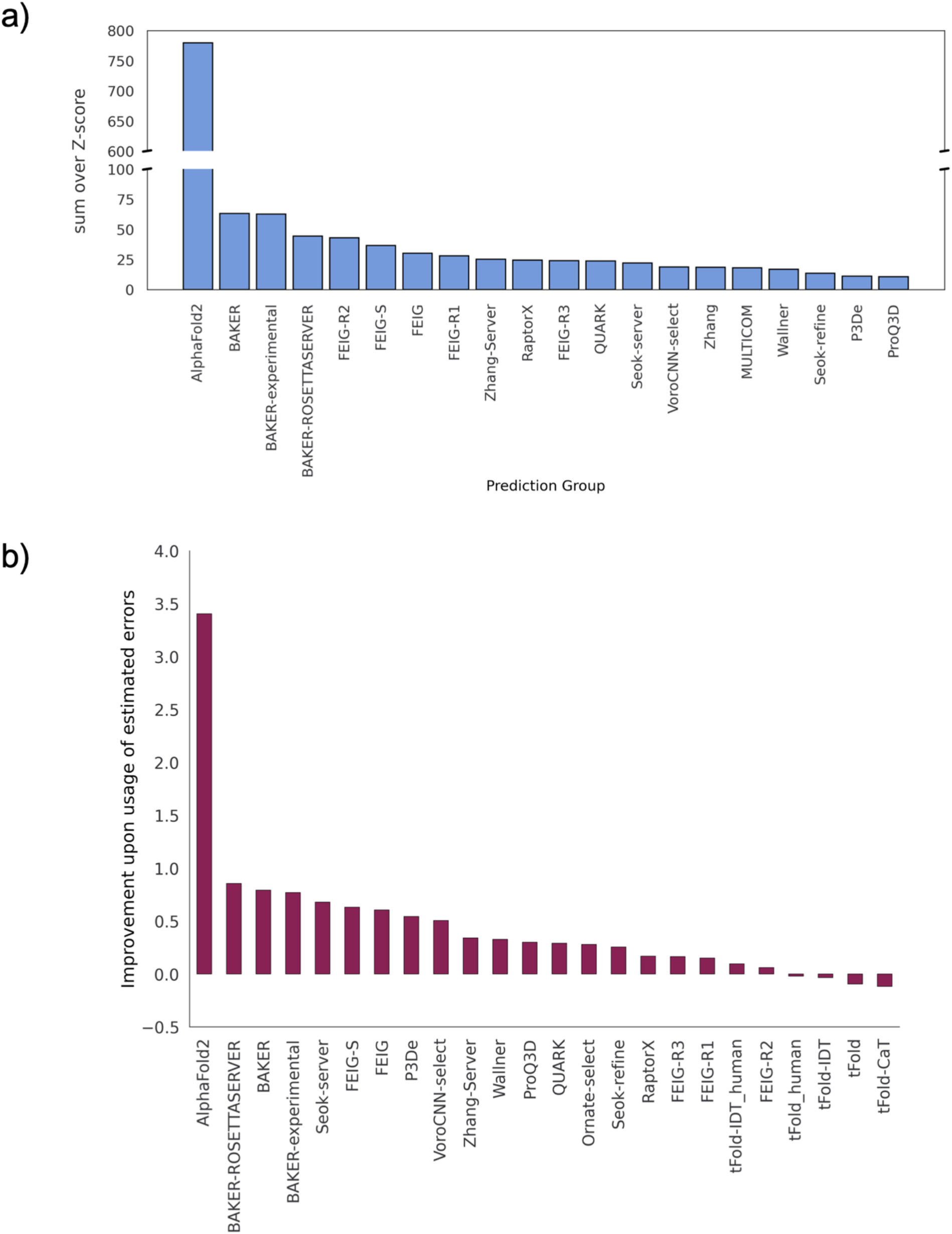
Ranking of predictions by reLLG. a) Group ranking by the reLLG ranking score for model #1 submissions. b) Improvement in performance for the top groups when the coordinate error estimates are used to weight the reLLG calculation. The 24 groups who were in the top 20 for either the weighted or unweighted scores are included.

We note that the top 3 groups are the same in this ranking as in the rankings using just targets for which diffraction data are available, but there are substantial differences in other methods near the top. Based on the comparisons discussed above, we believe that these differences reflect sampling error rather than a systematic difference between targets with and without diffraction data. Such sampling error should be reduced for the larger set of targets, further supporting the decision to use the reLLG Z-score as the primary ranking measure in this work.

#### Utility of coordinate error estimates in MR calculations

CASP participants are asked to contribute error estimates for their predicted models in the B-factor field of submitted PDB files. While the group ranking analysis in this study has been done using the information from those estimates, we also computed the reLLG scores substituting those estimates by a constant value. We then computed the difference between the sum of the reLLG scores for each group, either using or not using the error estimates. As can be observed in Fig. 5b, the general trend for the top scoring groups is that the inclusion of the error estimation in the reLLG calculation improves the score.

#### Accuracy self-assessment in the prediction category

The ability of the groups to identify their best models and rank them is an important aspect for prospective users, as many users will focus on the top model. Arguably, this is somewhat less important for MR models, as it is reasonably common (though not universal) to test a number of alternative models. One metric that can be used to score the accuracy of self-ranking is a rank correlation. We chose instead to use the fraction of the time that the #1 model is also the best of the 5 models submitted, because it is easy to understand and corresponds to one of the possible MR scenarios where only the best model is tested.

A scatter plot comparing the percentage of #1 models ranked correctly with the reLLG ranking score (Fig. 6a) shows that there is no overall correlation (correlation coefficient of −0.02) between the ability of an algorithm to predict structure and the ability to rank a set of predictions. This is unexpected, as one would expect ranking to be an essential component of successful prediction. Nonetheless, Fig. 6b shows that the most successful groups do better than random, with BAKER and FEIG-R1 doing best.

**Figure 6.**
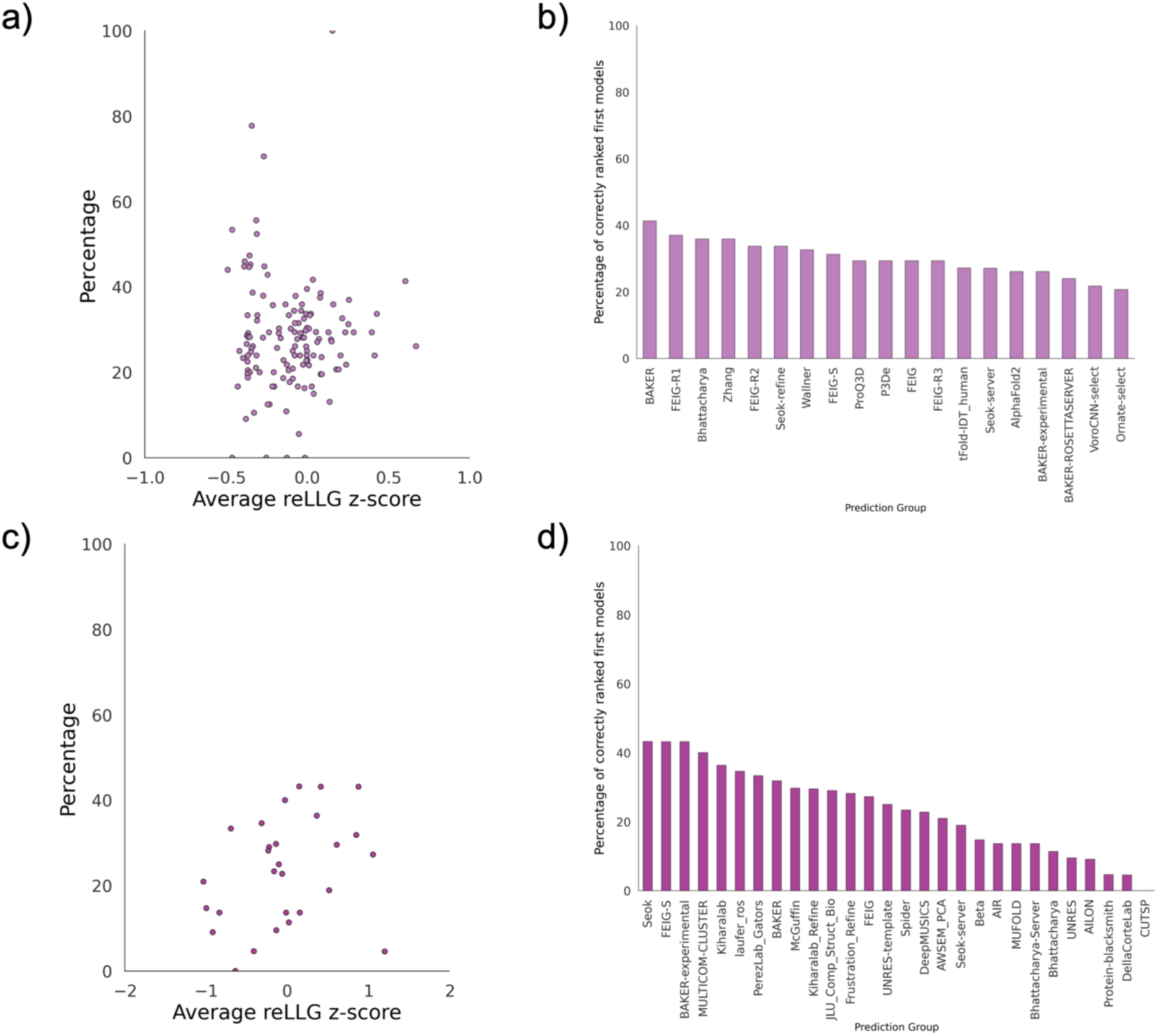
Percentage of targets for each group for which model #1 was the highest scoring in reLLG. Only the targets for which 5 models were submitted were considered. a) Scatter plot of percentage correct vs average reLLG Z-score for the prediction category. All groups are included except AlphaFold2, for which the average reLLG Z-score is 8.28 and the percentage correctly ranked is 26.1. b) Bar plot of percentage correctly ranked with the top 20 best groups from the overall prediction category ranking. c) Scatter plot for the refinement category, as in (a), including all groups. d) Bar plot, as in (b), with all groups from the refinement category.

### Refinement assessment

#### Refinement group ranking by reLLG scores

In this category, 36 groups contributed to 44 targets. Group rankings were computed in the same way as for the prediction round. To assess whether starting models were generally improved or degraded by refinement, we included in the ranking calculations a “naïve predictor”, who returns the starting model unchanged. One complication in scoring the naïve predictor was that the B-factor field of the PDB files containing the starting models did not typically contain estimated RMS coordinate errors; for consistency we evaluated the starting models by computing the reLLG score with all B-factors set to a constant. For this reason, even if a refinement group had left the coordinates unchanged but provided useful error estimates, they would have surpassed the naïve predictor.

Fig. 7a shows that the refinement of starting models is a difficult problem, as only 6 groups managed to consistently improve the models. In keeping with findings from other CASP metrics (Simpkin et al., this volume), the top 3 groups (FEIG, FEIG-S and DellaCorteLab) employed restrained molecular dynamics methods.

**Figure 7.**
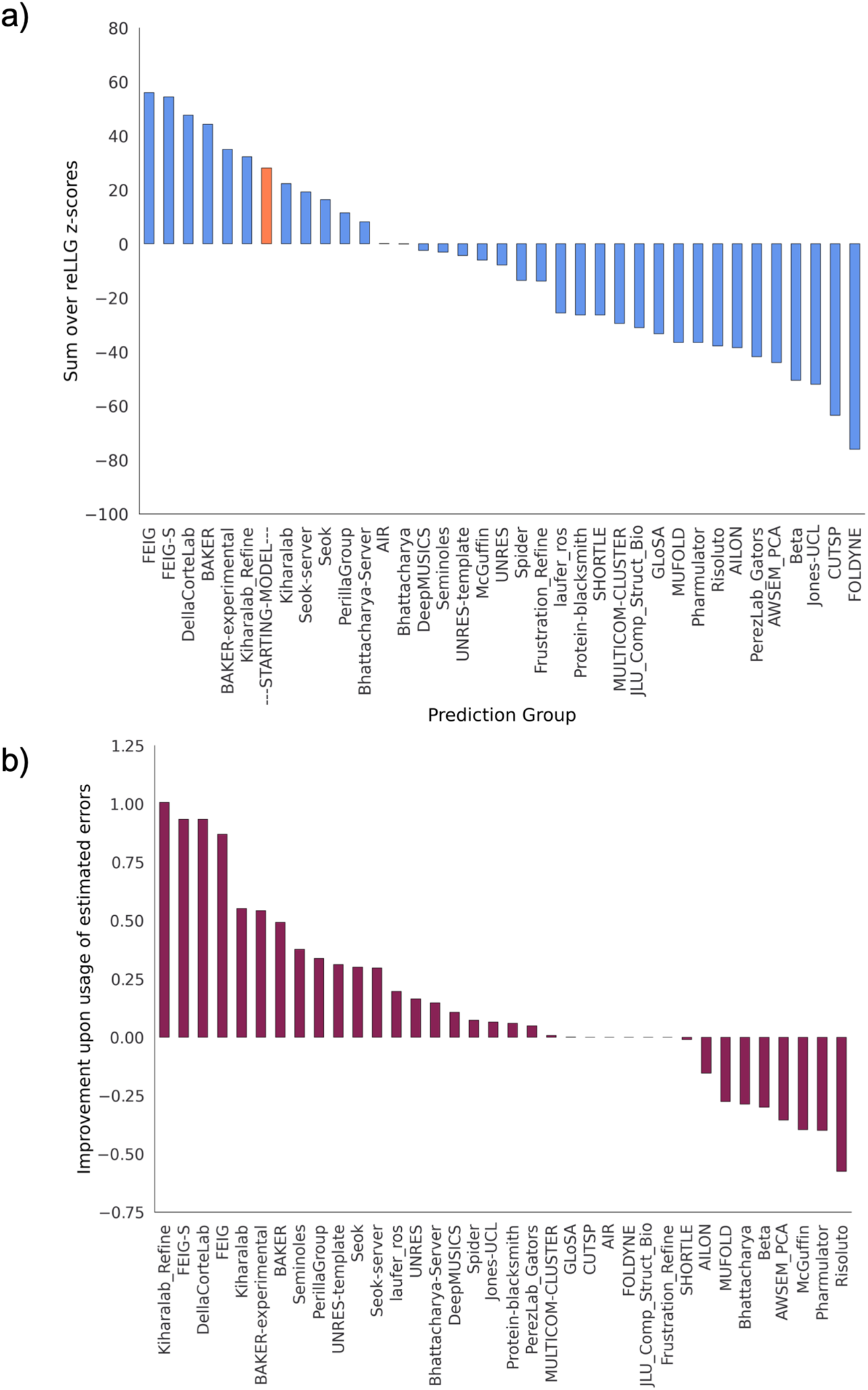
Ranking of refinement models by reLLG. a) Group ranking by the reLLG ranking score for model #1 submissions. b) Improvement in performance when the coordinate error estimates are used to weight the reLLG calculation.

#### Utility of coordinate error estimates

The effect of including the coordinate error estimates in the reLLG scoring was evaluated as for the prediction category. Fig. 7b shows that, again, considerable value was added to the model by including good coordinate error estimates. How much this added can be seen from an alternative ranking based on reLLG Z-scores computed with constant B-factors (Fig. S1), which therefore judge purely coordinate accuracy and not the accuracy of the error estimates. A comparison of Fig. S1 with Fig. 7a shows that only 3 groups outperform the naïve predictor, based only on coordinate accuracy: BAKER, FEIG and FEIG-S. The inclusion of error weighting moves Kiharalab_Refine up from 14^th^ position to 6^th^, above the naïve predictor, showing the real-world value of their excellent performance in coordinate error prediction, illustrated by Fig. 7b.

#### Accuracy self-assessment in the refinement category

There is a weak positive correlation (correlation coefficient of 0.31) between the ranking scores for different groups and their ability to correctly rank their best model as #1 (Figs. 6c and 6d). One would expect this to be a strength in deciding whether a starting model had been improved, but it is difficult to see why this ability should be more important for refinement than for the initial prediction where no overall correlation was seen.

#### Success of the refined models in MR

We performed MR using search models generated in the refinement category for those cases where diffraction data were made available. There were 13 targets that fulfilled this requirement. Four of these included extended submissions benefitting from 6 weeks of refinement in addition to the standard 3-week refinement submissions (T1034, T1056, T1067 and T1074). Further to this, T1053, T1067 and T1074 were double-barrelled cases with refinement performed on two initial starting models. In each of these cases one of the starting models was an AlphaFold2 prediction. This gave a total of 20 sets of refined models to be tested in MR. Refined models from 36 different groups were included with each group producing up to five models per target. Starting models were also used in MR for comparison. The full set of target details is provided in Table 1.

**Table 1.**
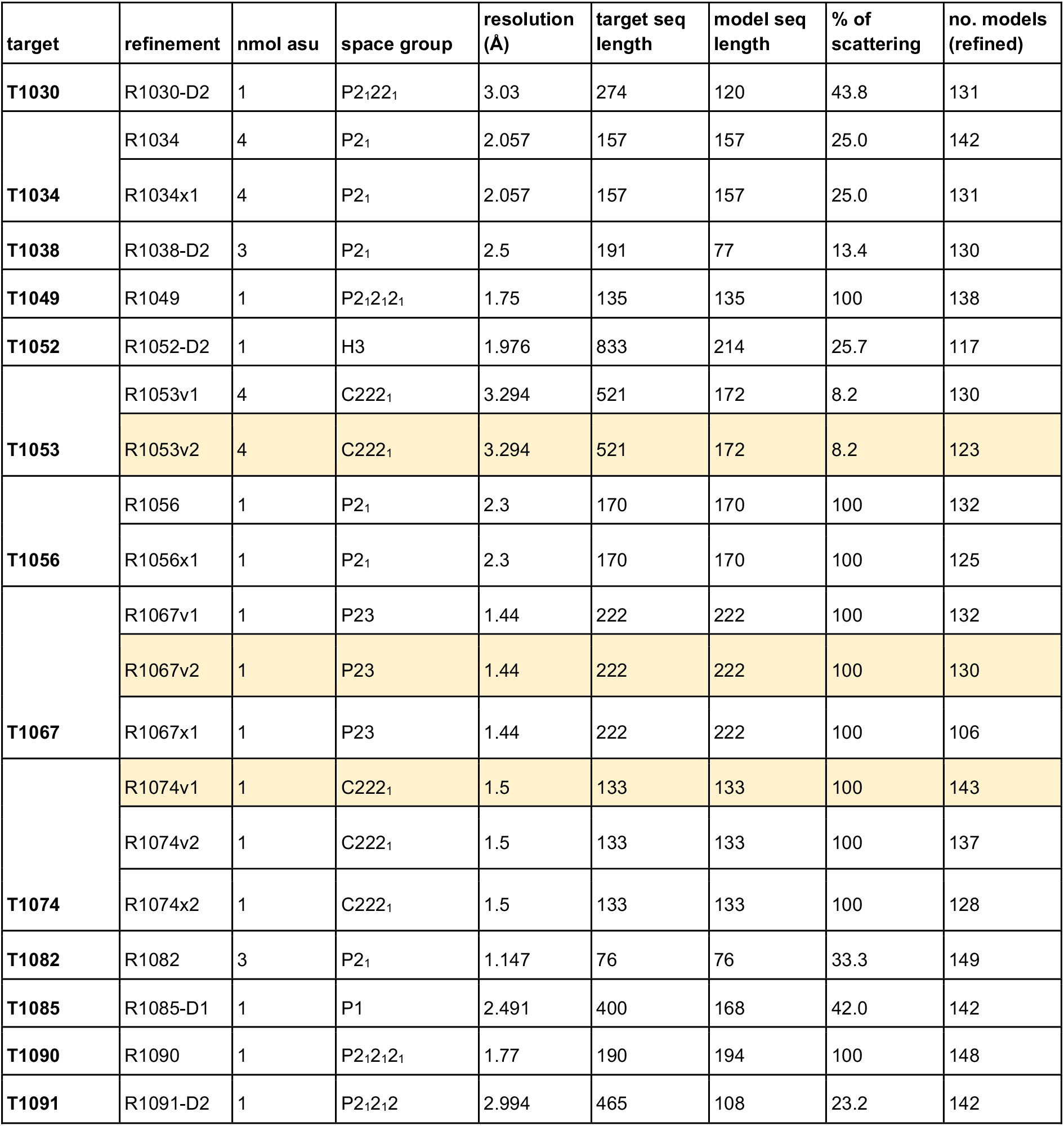
The set of prediction targets used in the refinement category where experimental diffraction data were available. The three double-barrelled cases had an additional refinement using an Alphafold2 starting model (highlighted). Refinements denoted with an “x” are where the model was refined for an additional 6 weeks. Cases with “D” denote starting models representing a single domain from the target.

The MrBUMP automated pipeline^29^ from the CCP4 suite^30^ (version 7.1.013) was used to take each set of refined search models and provide them to *Phaser*^31^ (version 2.8.3) to perform the MR trials. Some of the target crystal structures contained more than one protein molecule in the asymmetric unit, but we searched for only one copy to reduce the time taken for the MR run. For proteins with multiple components this is a more demanding test, because the signal in the MR search has a quadratic dependence on the fraction of the scattering accounted for by the model^5^. We deemed this to be an acceptable compromise as correct placement of the first copy is often indicative of a good chance of success in MR. The likelihood target in *Phaser* requires an estimate of the effective RMS coordinate error for the search model, which we set to 1.2 Å for all search models. For all of the refined models used, the B-factor field of the coordinate file was interpreted as an estimated RMS error, as discussed in Materials and Methods. To test if the solution in each trial was correct, we used phenix.famos from Phenix^28^ to calculate a mean log absolute deviation (MLAD) between the solved structure and the placed search model, accounting for any origin shift. A value of less than 1.5 for MLAD was used as the criterion for successful placement in MR.

Fig. 8 shows the overall performance of all the groups for each of the refined model sets. Of the 16 starting models for the 13 targets, only 5 of these proved to be successful search models in MR. Three of these were the AlphaFold2 predictions, with the remaining two being the starting models for R1034/R1034×1 (provided by the Seok server) and R1056/R1056×1 (from UOSHAN). Using these starting models, most groups that participated produced refined models that could also be used successfully in MR. In 9 of the remaining 13 cases (including extended targets) refined models were produced that were sufficient for correct placement in MR. The BAKER and FEIG groups proved to be the most successful, yielding positive results in 13 and 12 cases respectively. Notably, the same 6 groups appear at the top of the actual MR test as those above the naïve predictor in the reLLG ranking (Fig. 7a); the groups that ranked below the naïve predictor provided very few models that succeeded in MR when the starting model failed.

**Figure 8.**
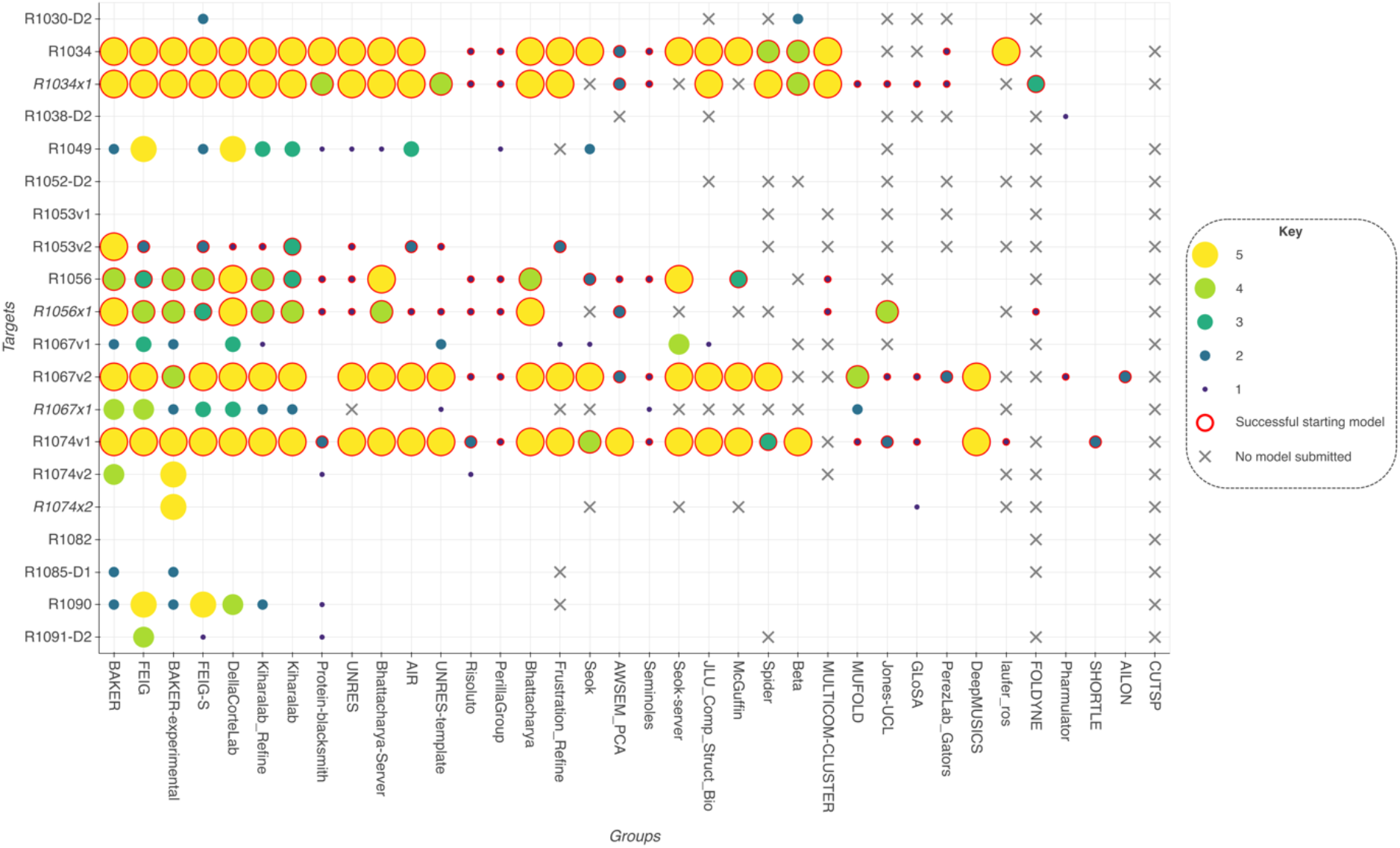
The plot shows the success of the group’s refined models in MR for each of the 20 refinement cases where experimental diffraction data were available. Groups are ordered from left to right by the number of cases where they produced at least one successful solution. Refinement cases involving the extended extra 6 weeks of refinement are shown in italics. The three cases where an AlphaFold2 prediction was used as the starting model are R1053v2, R1067v2 and R1074v1. Points are encircled in red where the starting model was also successful in MR.

An example of a successful refinement by the FEIG-S group of a starting model unsuitable as a search model in MR, for the target T1090, is shown in Fig. 9.

**Figure 9.**
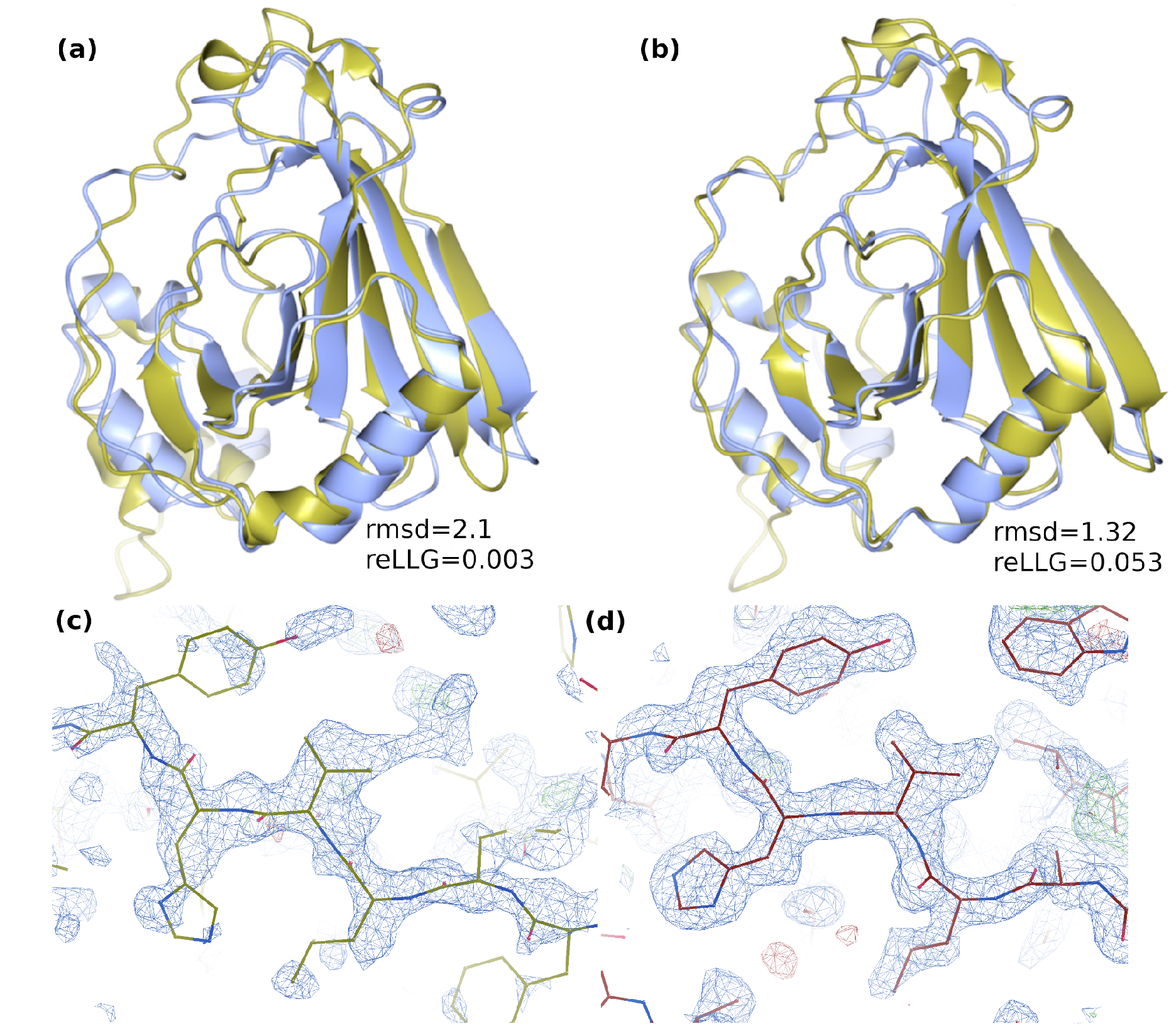
a) Starting model (gold) superimposed onto the target structure (blue). b) Refined model #1 from FEIG-S group (gold) superimposed onto the target (blue). Unlike the original refinement target (a), the FEIG-S refinement succeeded in MR and achieved an LLG of 145 and a local map correlation of 0.44. Panels (c) and (d) compare the quality of the map in the region around residue 153 from the phases generated from the placed refinement target (by superposition onto the placed FEIG-S model) and the MR-placed FEIG-S refinement. The phases generated by the model and the resulting electron density map are much improved by the refinement.

For two of the four targets subjected to the extra three weeks of refinement time, groups MUFOLD, Jones-UCL, GLoSA, FOLDYNE and UNRES-template were able to exploit the extra time to improve some of their models sufficiently to be suitable as search models for MR. Other groups including BAKER and FEIG were able to increase their success rate with the extra time. However, the overall results were mixed. For three of the four targets, the total number of models succeeding in MR declined following the extra refinement (Table 2). Only in the case of T1067 did numbers improve from 21 to 24. For the double-barrelled cases, it is clear that the high accuracy of the AlphaFold2 starting models made them very difficult to improve upon with refinement. Although the level of success for refined models produced from these was very high, the overwhelming majority of the models scored lower LLG and MLAD values in MR than the original AlphaFold2 predictions.

**Table 2.** Results for the “extended” models allowed an additional 6 weeks for refinement. The table shows the number of successes achieved in MR across all of the models for those groups that participated. The number of unique solutions at each stage of the refinement is also shown.

|  | 3 weeks |  | 6 weeks |  |
| --- | --- | --- | --- | --- |
| Target | models successful in MR | unique to 3 weeks | models successful in MR | unique to 6 weeks |
| **T1034** | 110 | 23 | 100 | 13 |
| **T1056** | 54 | 17 | 51 | 14 |
| **T1067** | 21 | 10 | 24 | 13 |
| **T1074** | 12 | 6 | 8 | 2 |

### Assessment of progress

As seen with many other CASP metrics, the quality of the AlphaFold2 models for MR represents a step change in what can be achieved. It is difficult to attach a numerical value to quantify progress in MR, but there is strong qualitative evidence. In previous rounds of CASP, the quality of models for MR was never measured for the most difficult (FM and FM/TBM) categories because almost none of the models were judged to bear sufficient resemblance to the targets to make that a meaningful exercise. In addition, this is the first occasion in which targets contributed to CASP were actually solved using submitted models.

Target difficulty in CASP is traditionally measured using a linear combination of target rankings by sequence coverage and sequence identity to the closest homologue of known structure^33^. Fig. 10a shows that model quality for MR, measured by the reLLG score, still depends on target difficulty, but there are useful models across the spectrum. In almost all cases, the best models are those produced by AlphaFold2. One striking example is their model #2 of T1078-D1, which achieves an reLLG score of 0.648, the highest seen for any of the targets; this is in spite of the difficulty level of this target, for which the best template in the PDB has a sequence identity of only 9.8% and a coverage of 71% of the target length. AlphaFold model #1 for T1053-D1 is a very good model for an even more difficult target, where the closest homologue (chain A of PDB entry 3akk^32^) has a sequence identity of only 7.2% and a coverage of 47.8%. Figs. 10b and 10c show the striking improvement over the best template. Where the AlphaFold2 algorithm still has difficulties, indicating room for improvement, can be seen from cases where low scores were obtained in spite of apparently modest difficulty levels; these are outlined in a dashed blue box at the bottom of Fig. 10a. The blue points in this box represent the best AlphaFold2 models for (from left to right) T1093-D2, T1100-D1, T1092-D1, T1083-D1, T1095-D1 and T1099-D1. These all represent cases of targets extracted from subunits of larger assemblies: T1083-D1 is a subunit of a homotetramer stabilized by coiled-coil interactions, T1092-D1, T1093-D2, and T1095-D1 correspond to three subunits of H1097, the phage AR9 RNA polymerase, T1099-D1 is a single subunit of the duck hepatitis B virus capsid and T1100-D1 is a subunit of a homodimer stabilized by a long coiled-coil interaction. Clearly the prediction of structure in the absence of the structural context is still a difficult problem. In spite of this, remarkably, the models of subunits of the phage AR9 RNA polymerase were good enough to play a pivotal role in solving the structure of this complex (Leiman *et al.*, this volume).

**Figure 10.**
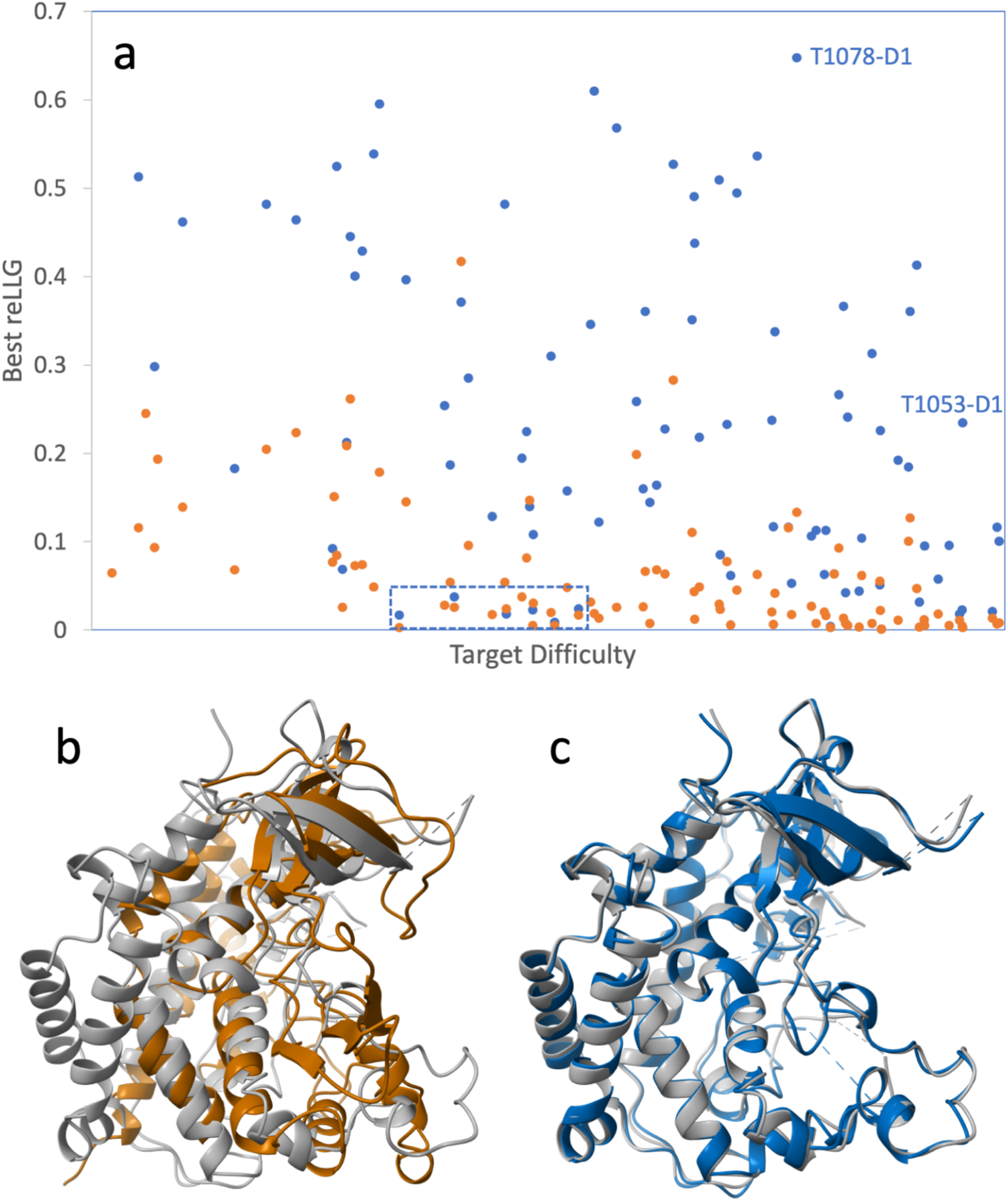
a) Model quality, measured by the reLLG score weighted by estimated RMS error, as a function of target difficulty. The points in blue represent the best AlphaFold2 model for each target, and the points in orange represent the best non-AlphaFold2 model for each target. b) Superposition of chain A of PDB entry 3akk^32^ (brown) on the structure of T1053-D1 (grey). c) Superposition of model #1 from AlphaFold (blue) on T1053-D1 (grey).

Methods of similar power to AlphaFold2, when they become readily available to the structural biology community, can be expected to play an increasing role in structure determination. We note that the development of the RoseTTAFold algorithm, inspired in part by features of AlphaFold2, has already enabled the determination of several structures that evaded previous efforts^34^.

## Discussion and Conclusions

Crystallographers have a great deal of experience carrying out MR with models derived from homologues with different levels of sequence identity. Although success in MR involves a combination of factors (quality and completeness of model, quality and resolution of diffraction data), a commonly-used rule of thumb is that MR is likely to succeed if there is a homologue with at least 30% sequence identity (for 100% sequence coverage). It is useful to also relate the reLLG directly to solvability.

Sequence identity correlates with solvability because there is a relationship between sequence identity and the effective RMS error, termed the VRMS in *Phaser*, which is an important parameter in LLG calculations in the MR search. The VRMS can be estimated from sequence identity, taking into account perturbations introduced by molecule size^18^. For a complete model of a 175-residue protein (a typical globular protein/domain size) with 100% identity to the target, the formula yields a VRMS of approximately 0.4 Å, the value assumed for the ideally imperfect model in the reLLG calculation (reLLG=1.0). As sequence identity degrades, the VRMS increases as predicted by the Oeffner et al. formula^18^, and this can be translated into a reduced reLLG, as shown in Fig. 11. A sequence identity of 30% thus translates into an reLLG value of slightly less than 0.1. The majority of AlphaFold2 structures across the difficulty scale reach this value, as well as a substantial fraction of the best models from other groups (Fig. 10a).

**Figure 11.**
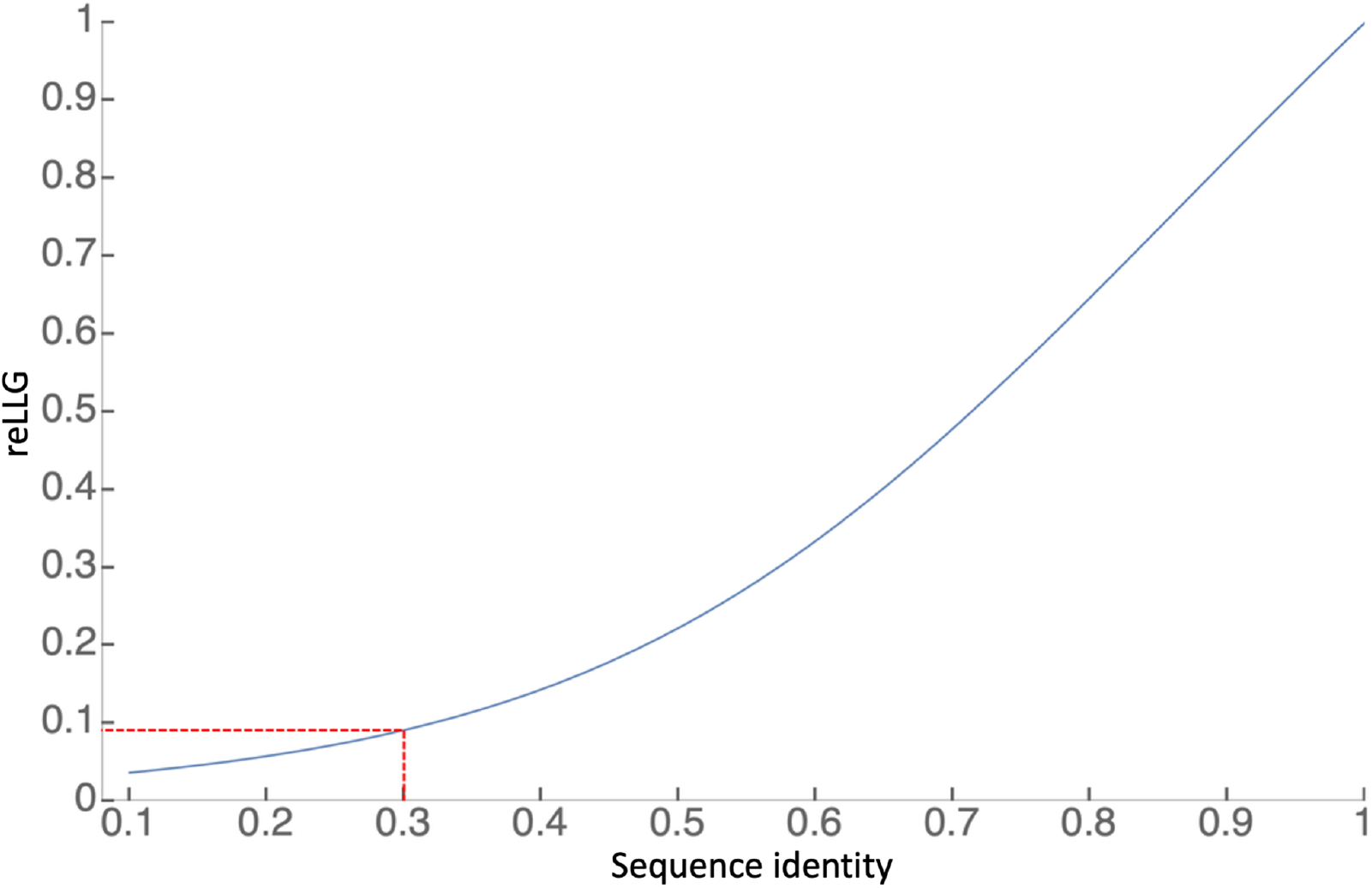
Translation of fractional sequence identity of a 175-residue protein into an equivalent reLLG value, assuming that the coordinate errors are all drawn from the same 3D Gaussian distribution inferred from the sequence identity. The dashed red lines show that a sequence identity of 0.3 would translate into an reLLG of about 0.091.

In this work we have not validated whether or not a reLLG over 0.1 is sufficient to solve the structures for which diffraction data are available. However, MR trials have been carried out as part of the high-accuracy assessment (Pereira *et al.*, this volume), addressing 32 targets and solving 30. Twenty-six of those required no editing in the Ample truncation procedure^4^, while a further three succeeded with truncated search models automatically generated by consideration of predicted residue errors. One required manual splitting of model domains. In a separate study^35^, in depth MR trials using only the AlphaFold2 models were carried out when data had become available for 34 crystal structures, from which 72 EUs had been defined. Of the 34 structures, 31 could be solved with AlphaFold2 models, 2 could be solved partially and one could not (though it could be solved with generic helix models), at least confirming the result for AlphaFold2 models.

### Relevance of refinement category in CASP

The CASP refinement category was instigated to encourage the development (and allow the evaluation) of expensive computational methods, ones for which most groups do not have the resources to apply to the large number of targets in the prediction round. In this category, a number of server-generated models are traditionally provided for further improvement. In CASP14, this pool of models was supplemented with 7 (non-server) AlphaFold2 models. We have seen that the best groups were consistently able to improve the server-generated refinement targets, but that most refinement methods degrade the AlphaFold2 models, as seen here for MR as well as for other CASP assessment measures (Simpkin *et al..*, this volume). The AlphaFold2 prediction algorithm (and the resources behind it) is better than even the compute-intensive refinement algorithms of other groups. This result is shown in Fig. 12 where, with one marginal exception (a slight improvement on an AlphaFold2 starting model), the AlphaFold2 model would have scored equal or higher on the reLLG score compared to the best refined model, even including the double-barrelled targets starting from AlphaFold2 models. If the initial AlphaFold2 predicted models had simply been resubmitted for each refinement target then AlphaFold2 would have topped the refinement rankings as well. In light of the highest quality predictions, the refinement category as it currently stands appears to have become redundant. Some consideration of potential future changes can be found elsewhere in this issue (Simpkin *et al..*, this volume).

**Figure 12.**
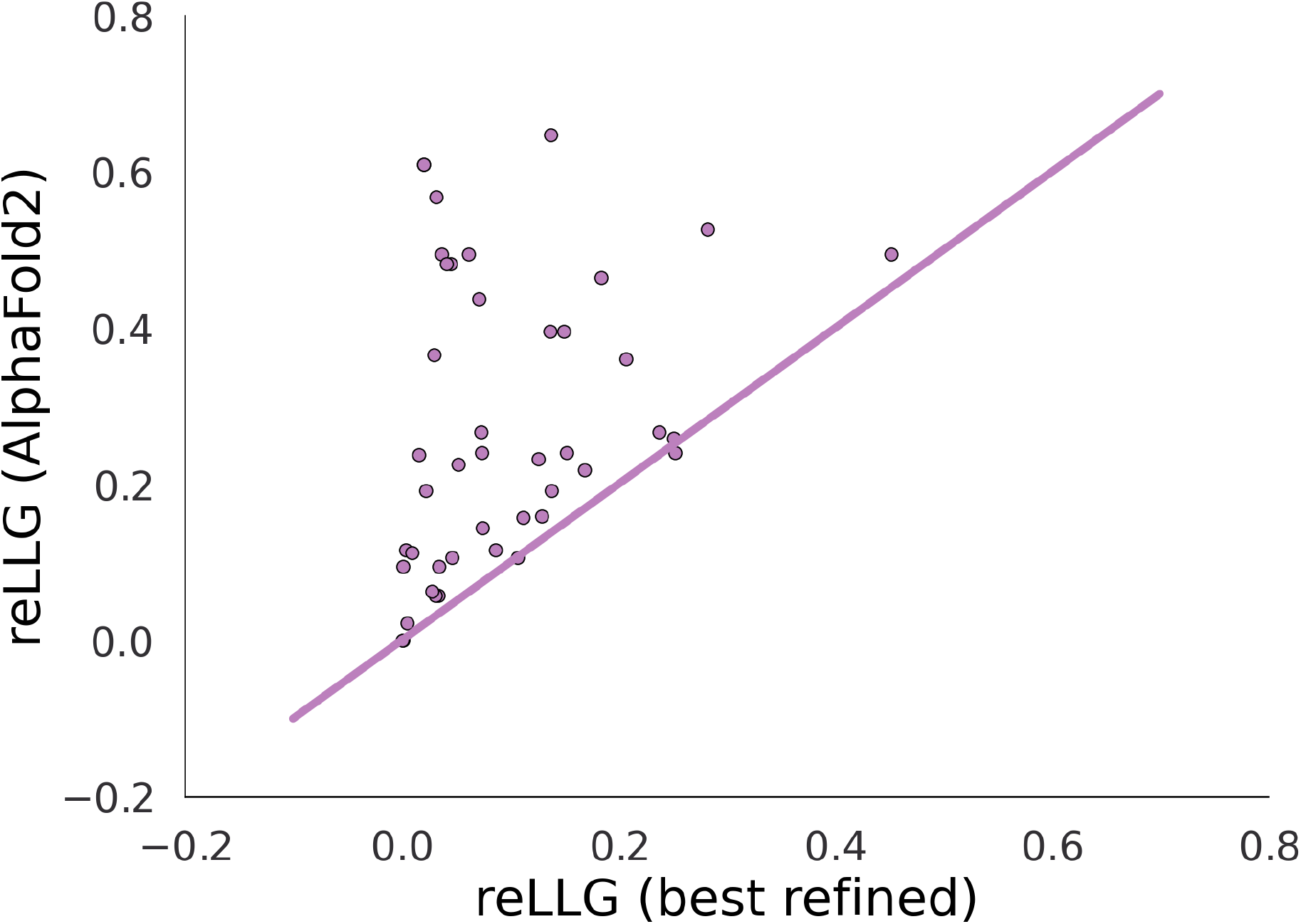
Scatter plot comparing reLLG scores of best AlphaFold2 model from the prediction round with the best model from the refinement round for each refinement target.

In conclusion, we have shown that the reLLG is a useful addition to the assessment metrics for CASP and should replace the metrics based on dLLG used in previous rounds. Although developed in the context of MR, it can be evaluated for models of structures determined by NMR, cryo-EM or with other structural restraints. It has a broad advantage over other metrics by combining assessment of coordinate accuracy with the assessment of the accuracy of the estimates of RMSD in coordinates. To further improve the reLLG of predicted models (and thus their utility in MR), groups should target estimates of individual atomic accuracy rather than grouped residue accuracy. It should also be understood by predictors that optimization of the reLLG ideally requires optimization as well of the predicted atomic B-factors, where the B-factor includes spatial and temporal sampling of coordinate locations in a dynamic molecular structure during the course of the structure determination data collection. To optimize reLLG in the current format, where there is one B-factor field in the submitted PDB file, would actually require submitting “error” values that, when translated into B-factors, produce the sum of the actual B-factor of the atom and the B-factor corresponding to the coordinate error. Since this would conflate more than one phenomenon in one number, CASP should facilitate the submission of different types of estimates for different purposes, by replacing the current PDB submission format with the flexible and extensible mmCIF format.

## Acknowledgements

We thank the experimentalists who provided diffraction data for a number of targets, and Andriy Kryshtafovych for the invaluable resources at the Prediction Center. This research was supported by a Principal Research Fellowship from the Wellcome Trust, awarded to R.J.R. (grant number 209407/Z/17/Z), the Biotechnology and Biological Sciences Research Council (grant BB/S007105/1), CCP4, and by institutional funds of the Max Planck Society to A.N.L. The authors have no conflict of interest to declare.

## Supplementary Material

**Figure S1.**
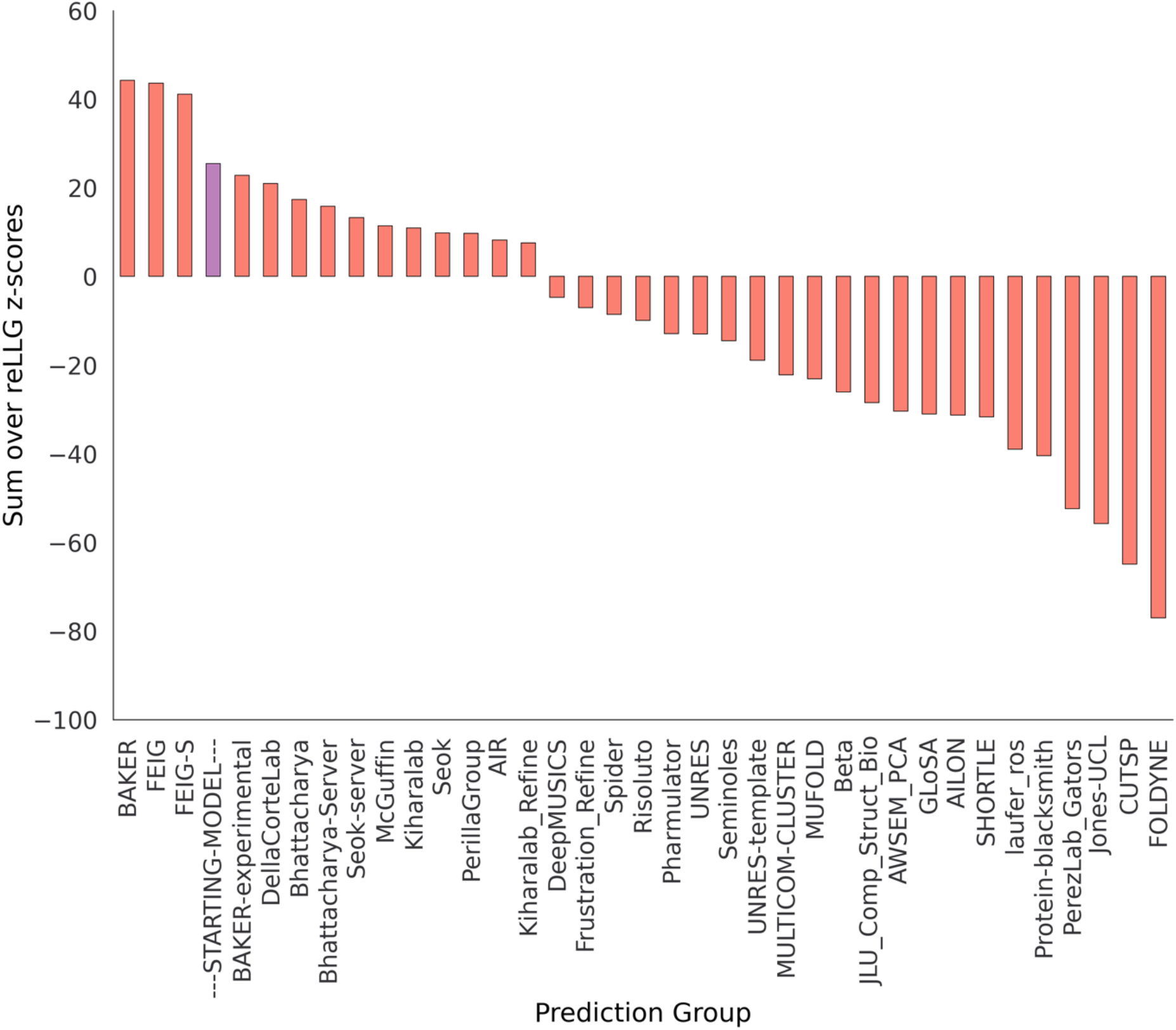
Alternative ranking of refinement groups by reLLG Z-score computed with constant B-factors. By this ranking, which focuses only on coordinate accuracy, only 3 groups outperform the starting model, which was also scored using constant B-factors.

**Figure S2.**
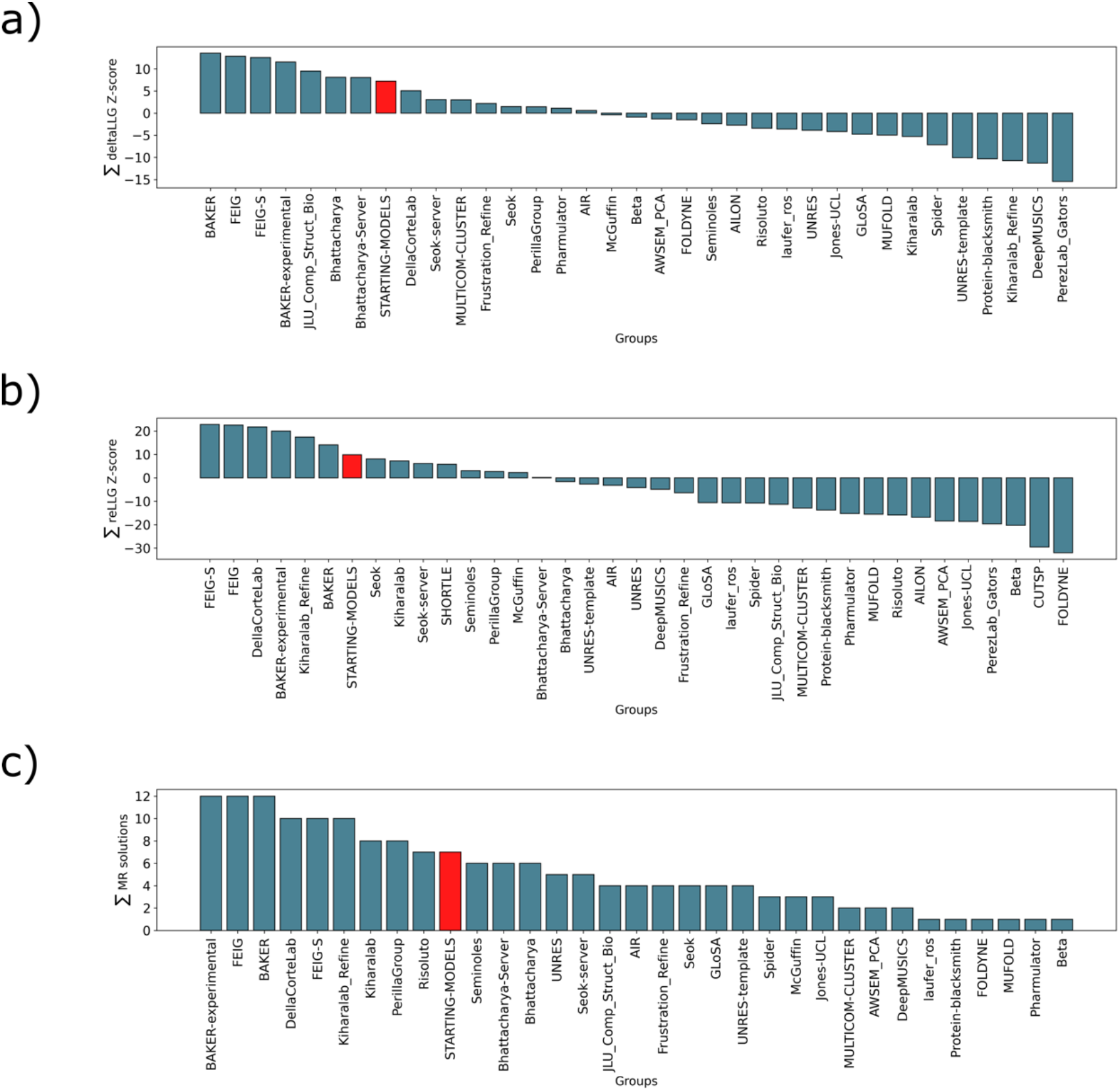
A comparison of alternative ranking strategies for the refinement methods against the 20 targets assessed with MR. These are ranked on the sum of dLLG Z-scores (a), the sum of reLLG Z-scores (b) and the sum of total solutions in MR (c). The scores were calculated using only model 1 and the naïve predictor is shown in red.

